# Spatial integration of sensory input and motor output in *Pseudomonas aeruginosa* chemotaxis through colocalized distribution

**DOI:** 10.1101/2024.03.13.584826

**Authors:** Zhengyu Wu, Maojin Tian, Sanyuan Fu, Min Chen, Rongjing Zhang, Junhua Yuan

## Abstract

The opportunistic pathogen *Pseudomonas aeruginosa* serves as a model organism for studying multiple signal transduction pathways. The chemoreceptor cluster, a core component of the chemotaxis pathway, is assembled from hundreds of proteins. The unipolar distribution of receptor clusters has long been recognized, yet the precise mechanism governing their assembly remains elusive. Here, we directly observed the relative positions of the flagellar motor and chemoreceptor cluster using flagellar filament labeling and gene editing techniques. Surprisingly, we found that both are located at the same cell pole, with the distribution pattern controlled by the polar anchor protein FlhF. Additionally, the efficient assembly of the chemoreceptor cluster is partially dependent on the integrity of the motor structure. Furthermore, we discovered that overexpression of the chemotaxis regulatory protein CheY leads to high intracellular levels of the second messenger c-di-GMP, triggering cell aggregation. Therefore, the colocalization of the chemoreceptor cluster and flagellum in *P. aeruginosa* serves to avoid cross-pathway signaling interference, enabling cells to conduct various physiological activities in an orderly manner.

## Introduction

In biological systems, the process by which proteins self-assemble into organized complex structures is widespread^1,2^. Pole-to-pole protein oscillations in the *Min* system ensure that the FtsZ ring, a crucial component of cell division, is placed precisely in the middle of the cell body^3,4^. In response to antibiotic treatment and heat stress, some bacteria generate multiple protein foci throughout the cytoplasm^5,6^. The polar localization of the secretion system, such as the type [secretion system, is mediated by targeting proteins and potentially facilitates host-pathogen interactions^7,8^. Thus, spontaneous assembly is accompanied by the formation of spatial patterns that organize the intracellular environment.

To grow and colonize in various ecological niches, bacteria have evolved a mechanism for migrating towards more favorable environments. The locomotion organ (the flagellar motor) and the chemotaxis pathway play crucial roles in achieving this goal. The former provides physical drive, while the latter offers directional guidance. The initiation of cell reorientation is controlled by motor switching, which is governed by the chemotaxis two-component signaling pathway. This pathway comprises transmembrane chemoreceptors, a histidine kinase CheA, an adaptor protein CheW, a response regulator CheY, and two adaptation proteins, CheB and CheR. Receptors collaborate with CheA and CheW to form chemotaxis complex in a hexagonal pattern^9,10^. Upon phosphorylation of CheY by the kinase at the complex, it freely diffuses to the motor and binds to FliM, a component of the motor switch complex, to affect motor switching^11–13^. It is worth noting that there exists diversity among different bacterial species in the placement of flagellar motors^14^, whereas chemoreceptor clusters are typically found at one or both ends of the cell^15^. These two structural units are functionally connected. However, it remains unclear whether there is an interaction in their spatial distribution and what the specific regulatory mechanism might be.

*Pseudomonas aeruginosa*, a common human opportunistic pathogen, possesses four chemosensory pathways that perform distinct functions and are stimulated by signal binding to 26 chemoreceptors^16^. Among them, the proteins encoded by chemotaxis-related genes collectively constitute the F6 pathway. Previous studies have suggested that the receptors involved in this pathway localize to the old cell pole^17^, similar to the flagellum^18^, implying colocalization of the receptors and the flagellum at the same pole; however, direct evidence for this colocalization is still lacking. Furthermore, the polar anchor protein FlhF has been reported to guide the flagellum to grow at the cell pole^19,20^. However, the mechanism by which chemotaxis complex-related proteins aggregate and self-assemble at the cell pole is controversial. The distribution of the chemotaxis complex in the peritrichous bacterium *Escherichia coli* has been extensively studied and attributed to several mechanisms, including stochastic self-assembly^21,22^, membrane curvature sorting^23,24^, and inefficient clustering in the lateral region^25^. However, these mechanisms cannot explain the unipolar distribution pattern observed in *P. aeruginosa*. Unlike *E. coli*, specific genes responsible for the placement of the chemotaxis complex have been identified in several bacterial species. In *Caulobacter crescentus*, chemotaxis complex assemble at the new cell pole with the help of TipN and TipF proteins^26^. A tripartite ParC-ParP-CheA interaction network was reported to promote polar localization of chemotaxis complex in *Vibrio parahaemolyticus*^27,28^. However, as there are no related genes in *P. aeruginosa*, the mechanism of its chemotaxis complex distribution remains a mystery.

Here, we combined gene editing and in vivo fluorescence imaging of flagellar filaments to directly observe the distribution of chemotaxis complex and flagellar motors, proposing a cooperative construction model of chemotaxis network and flagella during the entire division cycle of *P. aeruginosa*. The core focus of this study is to clarify the regulatory mechanism of its chemotaxis complex distribution. We found a substantial association between the assembly of the flagellar motor and the chemotaxis complex. The assembly efficiency of the receptor complex is influenced by core flagellar motor components, particularly the C-ring and MS-ring structures, while its assembly site is also affected by the polar anchor protein FlhF. Furthermore, by introducing exogenously expressed CheY protein, we found that this triggers the expression of c-di-GMP at a high level. From this, we infer that the colocalization of the chemotaxis complex and the flagellum in *P. aeruginosa* avoids the cross-pathway interference of signaling molecules, thus providing a guarantee for the coexistence of multiple chemosensory pathways.

## Results

### Robust generation of daughter cell with both chemotaxis network and flagellar motor

CheY has been shown to colocalize with chemoreceptors^17,29^. To visualize the distribution of chemotaxis complex in cells, we fused the gene encoding the enhanced yellow fluorescent protein (eYFP) to the *cheY* gene in the *P. aeruginosa* chromosome (Fig. 1A). All genetic modifications were carried out at the native position on the chromosome, allowing the expression of chemotaxis proteins in precise stoichiometry from their natural promoters (Fig. 1B). The mutant strain with *cheY-eyfp* was able to moderately expand on soft agar plates^17^, proving that its chemotaxis behavior was not notably affected, so it was selected for subsequent experiments. As shown in Fig. 1C, CheY-eYFP was mainly located at the cell pole, suggesting that CheA, CheW and CheY in *P. aeruginosa* form a signal transduction complex that was primarily distributed at the cell pole, as described previously^17^. Representative large-field images containing many cells are shown in Fig. S1. Additionally, we constructed a plasmid expressing CheA-CFP and electroporated it into the *cheY-eyfp* strain. Fluorescence imaging revealed a high degree of spatial overlap between CheA-CFP and CheY-EYFP (Fig. S2), confirming that CheY-EYFP accurately marks the location of the chemoreceptor complex. From our measurements, nearly 90% (332/372) of cells contain obvious receptor clusters, and factors such as fluorescence bleaching may have caused the loss of fluorescent spots in some cells. In addition, we observed the presence of fluorescent spots at both ends of some cells with a large aspect ratio (probably approaching cell division), indicating that the newly generated progeny cells will have a complete chemotaxis network. Next, we sought to observe the distribution of the flagella in cells simultaneously to understand the motility of future progeny.

**Fig. 1.**
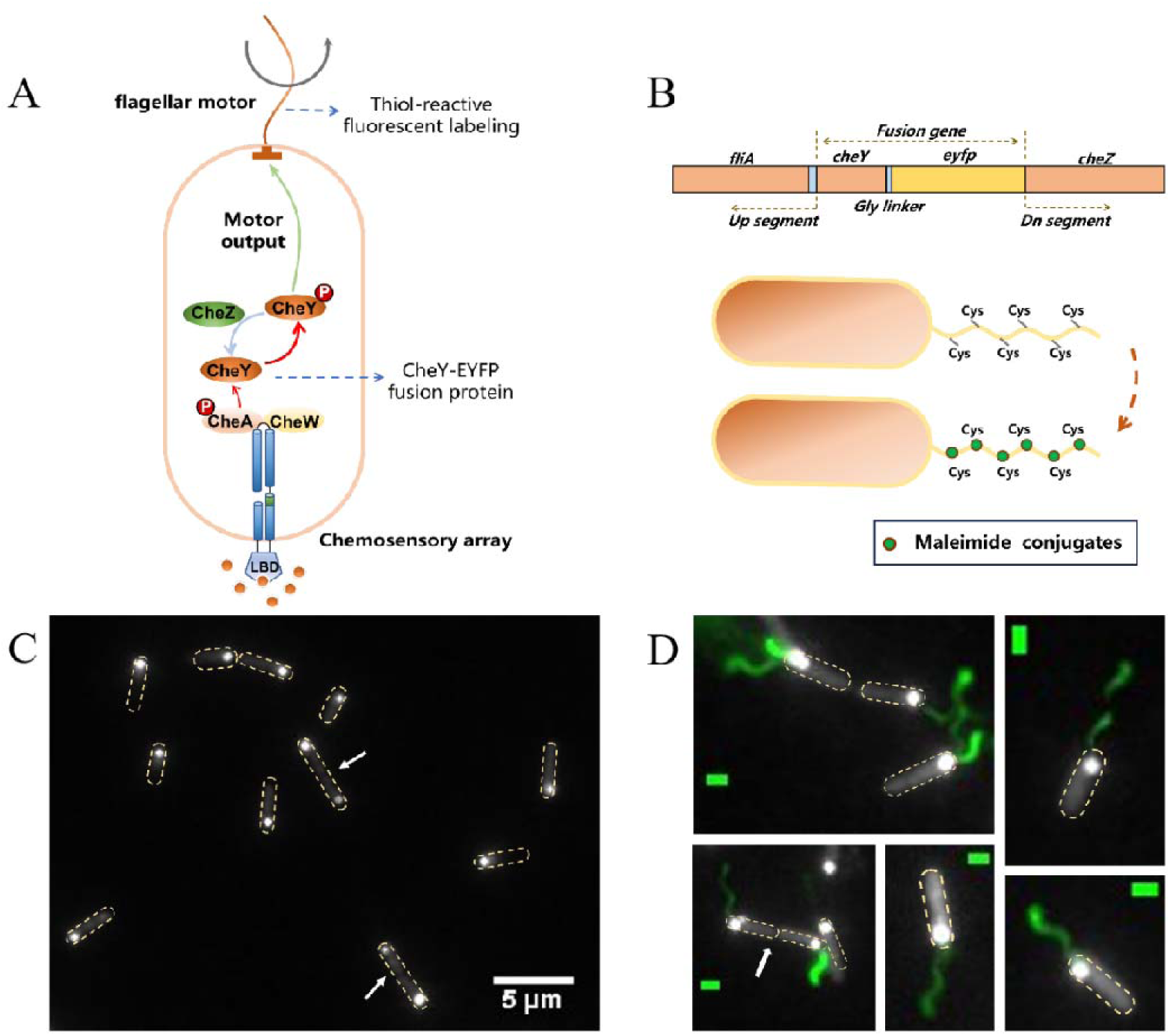
**A**. Schematic diagram of the chemotaxis signal transduction network and flagellar motor of *P. aeruginosa*. Fusion protein CheY-EYFP and fluorescently labeled flagellar filaments were used as markers to indicate the position of the chemotaxis complex and motor, respectively. **B.** The labeling mechanism of flagellar filaments and chemotaxis regulatory protein CheY. Filaments (with cysteine point mutation FliC^T394C^) were labeled through sulfhydryl-maleimide conjugation, and *cheY-eyfp* fusion with a 3× glycine linker was used to visualize chemotaxis complex positions. **C.** Localization of CheY-EYFP in the wild-type strain of *P. aeruginosa*. CheY-EYFP is mainly located at the single cell pole, and the white arrow points to individuals with obvious chemotaxis complex at both cell poles, which generally have a large aspect ratio of the cell body. The yellow dashed box marks the cell outline. **D.** The merged imaging of flagellar filaments and CheY-EYFP in the wild-type strain of *P. aeruginosa*, where flagellar motor and chemotaxis complex colocalize in cells. 145 cells with labeled flagella were observed, all of which exhibited consistent colocalization. White arrows point to individuals about to be divided, and the yellow dashed box marks the cell outline. The scale bar is 1 μm.

Our recent study demonstrated the successful labeling of the flagellar filament with thiol-reactive fluorescent dye by introducing cysteines into the flagellin FliC^30,31^ (Fig. 1B). To avoid fluorescence interference, we utilized a dye with an excitation peak near 568 nm, considering that the excitation peak of eYFP is 513 nm, and the emission peak is 527 nm. We employed a xenon lamp as the excitation light source and switched filters to enable simultaneous fluorescence imaging of flagellar filaments and chemotaxis complex within the same cell.

Remarkably, our findings revealed a surprising colocalization of chemotaxis complex and flagellar filaments at the same end of the cell body, and this colocalization was consistent, as shown in Fig. 1D. A typical large field was shown in Fig. S3. Similar to the chemotaxis complex, we observed the phenomenon of double flagella symbiosis in cells with a large aspect ratio, suggesting that *P. aeruginosa* has evolved a cell division mechanism with precise timing regulation. Before cell division, both poles of the mother cell assembled chemotaxis complex and flagellar motors. This ensures that daughter cells possess complete motility and chemotaxis, thereby greatly enhancing the environmental adaptability of the population.

### FlhF controls polar targeting but not flagellum-chemoreceptor colocalization

Despite potential differences in the physical and especially physiological environments at the two cell poles, it is unlikely that the unipolar distribution of the chemotaxis complex can be attributed to passive regulatory factors. The number and distribution of flagellar motors and chemotaxis complex vary among different bacterial species^14,15^, and the localization system for flagella has been well studied. Specifically, the FlhF-FlhG system, discovered in several monotrichous bacterial species, has been shown to control the location and number of flagella^32,33^. In *P. aeruginosa*, a knockout of *flhF* leads to mis-localized flagellar assembly^19,20^. Considering the consistent colocalization pattern between chemotaxis complex and flagellar motors in *P. aeruginosa*, we speculate that the distribution of chemotaxis complex is also regulated by similar molecular mechanisms.

To investigate the role of FlhF in the localization of chemotaxis complex, we constructed a Δ*flhF* strain for fluorescence observation. The results revealed that the chemotaxis complex no longer grow robustly at the cell pole (Fig. 2A), and the assembly positions of the flagellar motor change accordingly, colocalizing with the chemotaxis complex (Fig. 2B). Representative large fields containing many cells are shown in Fig. S1. Additionally, we categorized receptor cluster distribution patterns into three types: precise polar, near polar, and mid-cell. We artificially divided the intracellular area along the long axis of the cell. The two ends of the length accounted for 10% each, which we called the precise-polar domains, the middle 50% became the mid-cell domain, and the rest was called the near-polar domain. The type of complex distribution was determined by the partition where the center of the chemotactic complex fluorescent bright spot was located. The relevant statistics for the wild-type strain and the Δ*flhF* mutant are presented in Fig. S4, showing that the proportion of precise polar distribution of chemotactic complexes decreased by 12.8% after *flhF* knockout. We also quantified the proportion of individuals with obvious receptor clusters, which was more than 80% (182/221), similar to the wild-type strain. These experimental results suggest that the polar anchoring protein FlhF has a minimal impact on the assembly efficiency of *P. aeruginosa*’s chemotaxis complex, but it affects its unipolar distribution. Furthermore, a consistent colocalization between the flagellar motor and chemotaxis complex was observed, independent of FlhF. Thus, it is crucial to determine whether there is a causal relationship in the assembly order of these two structural units.

**Fig. 2.**
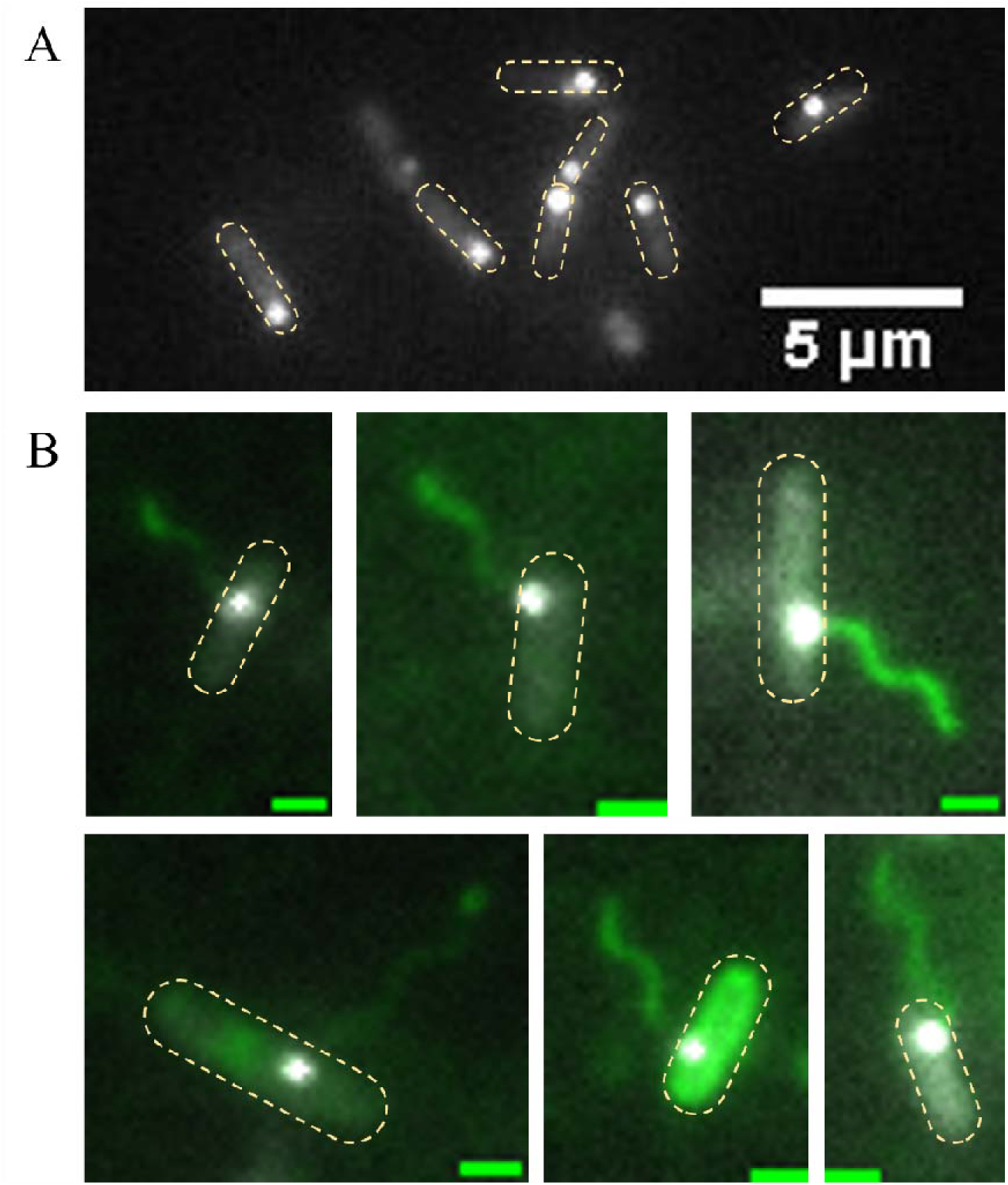
**A**. Localization of CheY-EYFP in the Δ*flhF* strain of *P. aeruginosa*. CheY-EYFP is no longer robustly distributed at the single cell pole. The yellow dashed box marks the cell outline. **B.** The merged imaging of flagellar filaments and CheY-EYFP in the Δ*flhF* strain of *P. aeruginosa*, flagellar motor and chemotaxis complex still colocalize in cells. 101 cells with labeled flagella were observed, all of which exhibited consistent colocalization. The yellow dashed box marks the cell outline. The scale bar is 1 μm.

### Core motor structures influence chemoreceptor complex assembly

Cryo-electron microscopy has successfully revealed the complete flagella structure in various bacterial species. This structure includes multiple components such as the inner-membrane MS ring, the cytoplasmic C ring, and the internal protein secretion system, all of which exhibit strong similarities^34^. This suggests that the core structure of the flagellar motor is highly conserved. The MS ring, composed of 26 FliF proteins, forms the foundation of the flagellar structure^35^. In addition to serving as a mounting platform for the C ring, the MS ring also acts as a protective shell for the internal protein secretion machinery^36^. FliG, the main component protein of the C ring, binds directly to the cytoplasmic surface of FliF in a 1:1 ratio^37^. Thus, the efficient assembly of the C ring requires preassembly of the MS ring. To observe the distribution of the chemotaxis complex following the disruption of the C ring and MS ring, we constructed Δ*fliG* and Δ*fliF* mutants.

We performed fluorescence observation on the Δ*fliG* mutant, which has a disrupted flagellar motor C ring. The results showed that the proportion of Δ*fliG* cells with obvious chemoreceptor clusters decreased significantly compared to the wild-type cells (59.8%, 140/234), although the chemotaxis complex remained at a single cell pole. We also examined the Δ*fliF* mutant, which has a disrupted flagellar motor MS ring. Similar to the Δ*fliG* mutant, the proportion of individuals with obvious chemotaxis complex decreased significantly (62.5%, 202/323). Subsequently, we constructed a Δ*flhF*Δ*fliF* mutant and observed that the proportion of individuals with obvious chemoreceptor clusters further decreased (50.7%, 341/672). To ascertain whether it is motor integrity rather than functionality that influences the efficiency of chemotaxis complex assembly, we constructed a stator mutant (Δ*motA*Δ*motCD*). In this mutant, the motor is completely stalled while the structure remains intact. We found that the mutant performed similarly to the wild-type strain in terms of chemotaxis complex assembly (84.3% of individuals with obvious clusters, 204/242). The fluorescence imaging of receptor clusters for multiple mutants in this section is shown in Figure. 3A, and corresponding large-field images are provided in Fig. S1. We utilized Western blotting to measure the expression levels of CheY, which were found to be similar across the different strains (Fig. 3B). These further substantiated that the observed phenomenon is based on the structural integrity of the motor rather than the protein expression level. In contrast, the *cheA* knockout strain was constructed to disrupt the assembly of the chemoreceptor complex. We fluorescently labeled its flagellar filaments and found that its phenotype was no different from that of the wild-type strain (Fig. S5), indicating that the assembly of the chemotactic receptor complex does not affect the flagellar assembly efficiency. The P ring (mainly composed of FlgI) is thought to act as a bushing for the peptidoglycan layer, and its absence results in partial assembly of the motor structure^38^. We constructed Δ*flgI* mutant and found that the proportion of cells with distinct chemotactic complexes was similar to that of the wild-type (Fig. S6). Additionally, we quantified the proportion of cells with receptor clusters (Fig. 3C). Overall, our findings suggest that the polar anchor protein FlhF and the structural integrity of the motor are crucial for the formation of chemotaxis complex. The former guides them to the appropriate site, while the latter influences the assembly efficiency of the chemoreceptor complex.

**Fig. 3.**
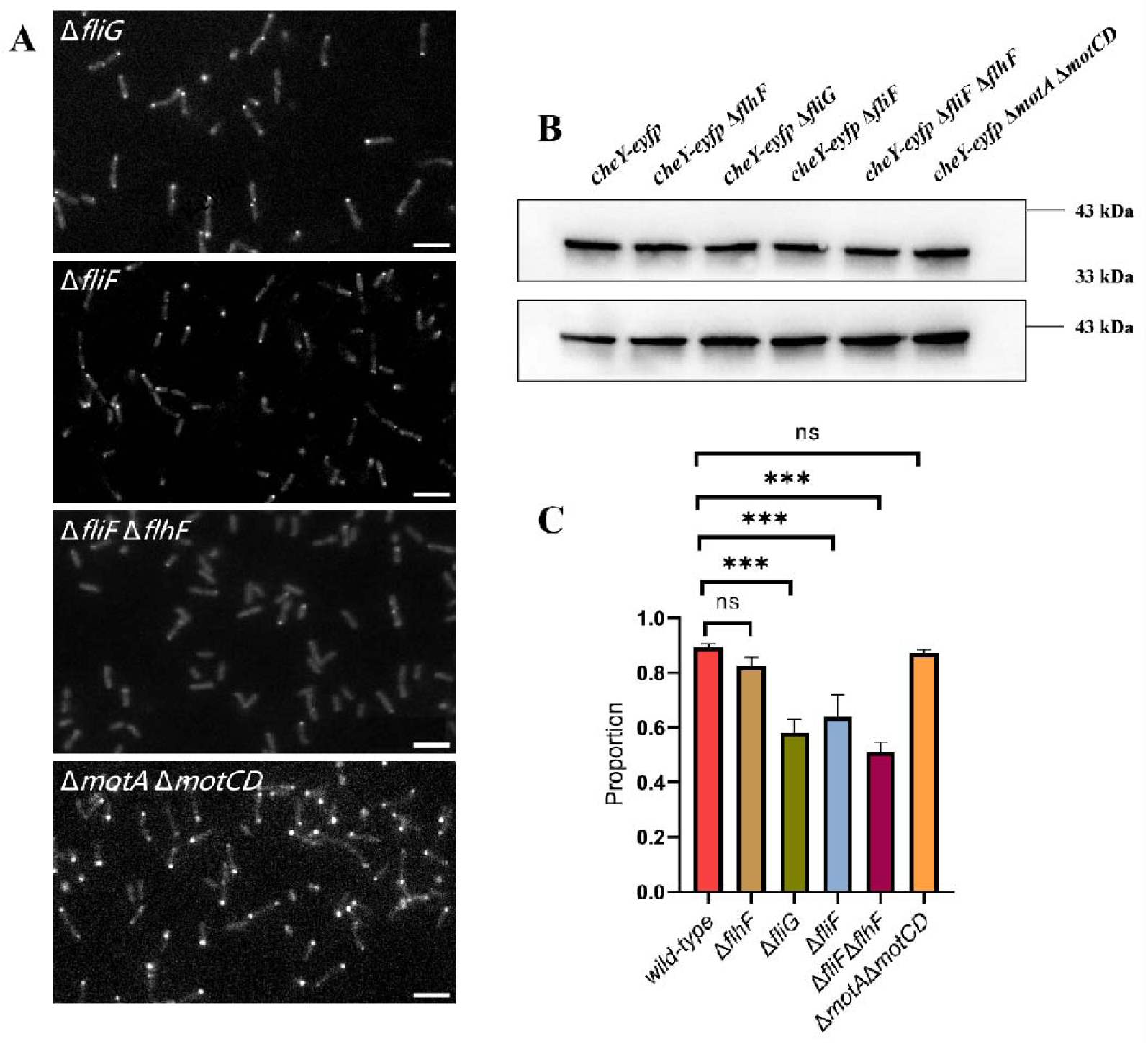
**A**. Localization of CheY-EYFP in various *P. aeruginosa* strains. The scale bar is 10 μm. **B.** Western blot analysis was performed to detect CheY expression in various *P. aeruginosa* strains. β-actin was used as the housekeeping protein. **C.** Occurrence probability of obvious chemotaxis complex in *P. aeruginosa* wild-type and several mutant strains. The proportion of individuals with obvious chemotaxis complex decreased significantly in the motor-incomplete strains (Δ*fliF* and Δ*fliG*), and this value was further reduced after *flhF* knockout (Δ*fliF*Δ*flhF*). The Δ*fliF* and Δ*motA*Δ*motCD* strains have a similar chemotaxis complex occurrence probability as the wild-type strain. The number of cells analyzed for each strain (from left to right) were: 372, 221, 234, 323, 672, and 242. ‘***’: significant difference (P-value < 0.0001), ‘ns.’: no significant difference (P-value > 0.05).

### Colocalization of chemotaxis complex and flagellar motor avoids cross-pathway interference in *P. aeruginosa* signal transduction

The distribution patterns of receptors associated with various signal transduction pathways in *P. aeruginosa* vary considerably. For instance, the biofilm formation-related receptor WspA, unlike the chemotaxis receptors discussed here, is distributed throughout the cell. This distribution is thought to enhance its sensitivity to mechanical perturbations of the cell membrane^39^. Previous studies speculated that the highly consistent position of chemotaxis complex and flagellar motors enhances bacterial chemotaxis performance^28^. However, the diffusion time of the phosphorylated chemotaxis regulatory protein CheY-P across the longest distance in a bacterial cell body (along the cell’s long axis) is approximately 100 ms^40,41^, whereas the timescale for the chemotaxis temporal comparison is on the order of seconds^42^. Additionally, a study by Fukuoka and colleagues reported that intracellular chemotaxis signal transduction requires approximately 240 ms beyond CheY or CheY-P diffusion time^41^. Moreover, the CCW/CW interval of the *P. aeruginosa* flagellar motor under normal conditions is 1-2 s, as determined by bead assays^30^ or tethered cell assays^43^. Taken together, these indicate that for *P. aeruginosa*, which moves via a run-reverse mode, the potential 100 ms reduction in response time due to co-localization of the chemotaxis complex and motor has a limited effect on overall chemotaxis timing. Consequently, the physiological significance of the colocalization of chemoreceptors and flagellar motors remains unclear.

*E. coli* mediates chemotaxis through a single pathway involving five types of chemoreceptors^44^. As a core regulatory protein for chemotaxis, CheY participates in multiple processes such as adaptation and chemotaxis response, with CheY-P binds to the motor to regulate switching^11^. In contrast, *P. aeruginosa* has four distinct chemosensory pathways that perform different functions, involving different CheY homologs, and are stimulated by signals binding to 26 types of chemoreceptors. Consequently, *P. aeruginosa* possesses more complex chemosensory pathways^16^. We thus hypothesized that the co-localization of chemoreceptors and flagellar motors in *P. aeruginosa* ensures locally distributed CheY-P molecules, thereby eliminating the need for a high level of intracellular CheY-P and avoiding potential side effects on other signaling pathways.

To test the potential effect of an increased intracellular CheY-P level, we constructed the CheY expression plasmid *cheY*-pJN105, transformed it into the wild-type *P. aeruginosa* strain, and induced it with varying concentrations of arabinose. We observed that the higher the inducer concentration, the more pronounced the cell aggregation became in the field of view (Fig. 4A). The transition from planktonic individuals to aggregated communities is similar to biofilm formation, which is known to be accompanied by a significant increase in intracellular c-di-GMP levels.

**Fig. 4.**
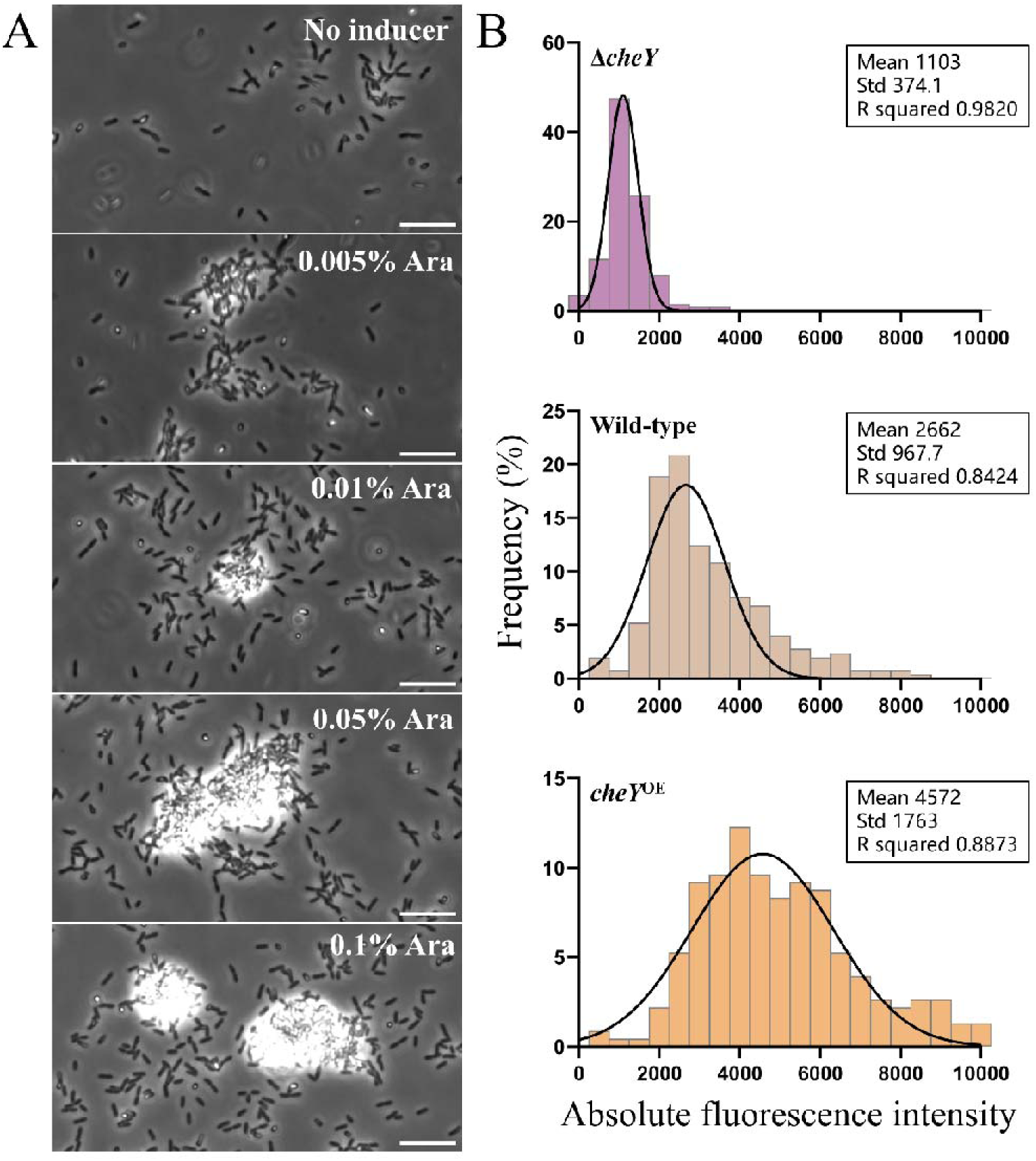
**A**. The evolution of cell aggregation as the intracellular CheY concentration increases by induction with higher concentrations of arabinose. The scale bar is 10 μm. **B.** Quantitative characterization of intracellular c-di-GMP levels at different CheY concentrations. From top to bottom, they correspond to the Δ*cheY* strain (N=198), wildtype strain (N=249) and CheY overexpression strain (N=228), respectively.

To investigate whether a higher level of intracellular CheY-P increased the intracellular c-di-GMP level, we sought to detect the c-di-GMP level. We introduced the plasmid pCdrA-gfp into *P. aeruginosa*, using it as a c-di-GMP biosensor with the fluorescence intensity proportional to the c-di-GMP level^45^. To ensure the coexistence of the plasmids, the intracellular CheY concentration was controlled by the plasmid *cheY*-pME6032. The intracellular CheY levels induced by varying arabinose concentrations were measured, showing clear differences across the concentration gradient (Fig. S7). We found that the average single-cell fluorescence intensity of the CheY overexpression strain was 71.8% higher than that of the wild-type strain, while the c-di-GMP level of the *cheY* deletion strain was 58.6% lower than that of the wild-type (Fig. 4B).

The colocalization of both the signaling source (the kinase) and sink (the phosphatase) at the chemoreceptor complex at the cell pole results in a rapid decay of CheY-P concentration within approximately 0.2 µm from the cell pole, leading to a nearly uniform distribution elsewhere in the cell, as demonstrated by Vaknin and Berg^46^. This spatial arrangement effectively confines high CheY-P levels to the pole region. When the motor is also localized at the cell pole, the need for elevated CheY-P concentrations throughout the cytoplasm is reduced. Therefore, the colocalization of chemotaxis complex and flagellar motor facilitates the precise regulation of cell swimming direction within the low intracellular CheY-P concentration threshold, using locally distributed CheY-P molecules. This helps to avoid the occurrence of the aforementioned cross-pathway interference. To further confirm that CheY overexpression promotes aggregation through increased c-di-GMP levels, we performed additional experiments co-overexpressing CheY and a phosphodiesterase (PDE) from *E. coli* to reduce intracellular c-di-GMP. These experiments showed that PDE expression mitigates cell aggregation caused by CheY overexpression in *P. aeruginosa* (Fig. S8).

## Summary and Discussion

*P. aeruginosa* harbors multiple signal transduction systems that regulate flagella-mediated swimming motility (Che pathway), pili-mediated interface twitching motility (Pil pathway), and biofilm formation (Wsp pathway)^16^. These systems allow it to thrive in various ecological niches within complex external environments. Here, employing a combination of chromosomal fluorescent protein fusion and flagellar labeling techniques in living cells, we directly observed that the chemotaxis complex of *P. aeruginosa* Che pathway and the flagellar motor share a high degree of consistency in their assembly sites. We found that the chemotaxis complex, comprising Che proteins, was consistently located at the cell pole where the flagellum was positioned. Based on these observations, we deduced the construction mode of the chemotactic network and flagellar motor during the entire cell growth cycle (Fig. 5A). As the cell body matures and elongates, a new flagellum grows at the opposing cell pole, accompanied by the assembly of a fresh chemotaxis complex. The cell then undergoes division from the middle, generating two daughter cells, each equipped with fully functional motility and chemotaxis capabilities. This ensures robust inheritance of these chemotaxis and motility-related macromolecular machines.

**Fig. 5.**
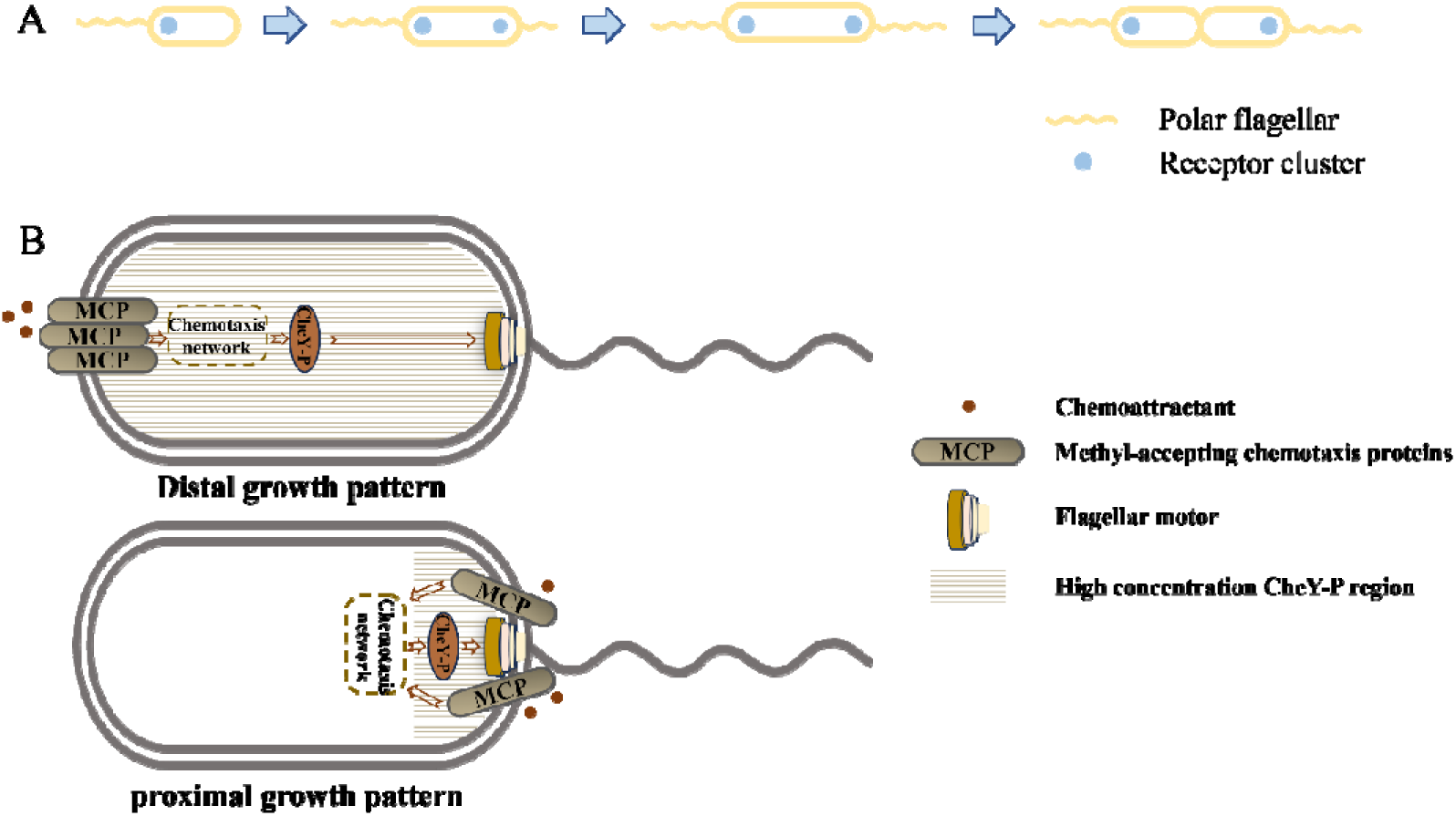
**A**. The construction mode of the chemotaxis network and flagellar motor throughout the complete cell growth cycle. **B**. The proximal growth mode of the *P. aeruginosa* flagellar motor and receptor clusters will effectively regulate the spatial range of CheY action, thus avoiding unintentional cross-pathway regulation.

Based on our understanding of the regulation mechanism of flagellar position in *P. aeruginosa*, we constructed a Δ*flhF* mutant and found that the position distribution of its chemotaxis complex is actively regulated by FlhF-mediated molecular mechanisms, rather than being passively restricted by membrane curvature factors as seen in *E. coli*. A similar active regulation phenomenon was previously reported in *Vibrio cholerae*, with the corresponding molecular regulation mechanism detailed extensively^28^. The flagella and chemotaxis complex of newborn *V. cholerae* cells are located at the old cell pole, with the polar anchor protein HubP distributed at both cell poles. As the cell grows, HubP recruits the ParA homologue ParC. Subsequently, ParC recruits ParP to assemble a new chemotaxis complex at the new cell pole. Upon cell division, HubP relocates to the middle of the cell, ensuring its robust presences at both poles of the daughter cells. Here, HubP recruits ParA homologues including FlhG, which influence flagellar motor assembly. However, previous experiments confirmed that the cell cycle-dependent polar localization of ParC and chemotactic proteins is independent of the flagellar position regulatory protein FlhF. This suggests that the localization of flagellar motors and chemotaxis complex in *V. cholerae* is controlled separately through different pathways. In contrast, flagellar motors were no longer robustly distributed at the cell pole after *flhF* was knocked out in *P. aeruginosa*, whereas the assembly site of chemotaxis complex and flagellar motor remained remarkably consistent. This suggests that FlhF regulates the distribution patterns of both and implies an interaction between them.

We further constructed Δ*fliG* and Δ*fliF* mutants, which disrupted the assembly of the flagellar motor. Surprisingly, we observed that the assembly of chemotaxis complex was impaired under these conditions, with a significant decrease in the proportion of individuals exhibiting obvious Che protein fluorescent bright spots. In contrast, the chemotaxis complex assembly in the stator mutant (Δ*motA*Δ*motCD*) was similar to that of the wild-type strain. It should be noted that *P. aeruginosa* has a four-tiered transcriptional regulatory circuit controlling flagellar biogenesis, in which FliG and FliF belong to class II and are regulated by the master transcriptional regulator FleQ^47^. Chemotaxis complex-related genes (*cheA*, *cheW*, etc.) are regulated by intracellular free FliA proteins. In *E. coli*, FliA is bound and inhibited by FlgM, and upon hook assembly, FlgM is secreted outside the cell, allowing FliA to trigger transcription of class III genes^48,49^, which include the chemosensory genes. This implies that if the hook is not assembled, late genes (including chemoreceptors) should not be expressed. However, Kaplan et al. reported that chemotaxis complexes were observed in *Shewanella oneidensis* Δ*fliF* mutant^50^, suggesting a different assembly process from that of *E. coli*. We therefore investigated whether knocking out *fliF* or f*liG* affects the expression level of *che* genes in *P. aeruginosa*. Since most experiments in this study focused on CheY, we performed Western blotting to measure CheY expression levels in wild-type, Δ*fliF* and Δ*fliG* strains, and found no significant differences (Fig. 3B). These results suggest that the observed decrease in the proportion of chemotaxis complexes after knocking out *fliF* and *fliG* should be attributed to the lack of motor assembly rather than changes in Che protein expression levels. To eliminate the potential confounding effects of FlhF and flagellar motors, we constructed a Δ*flhF*Δ*fliF* mutant strain. Under this condition, the proportion of individuals with obvious Che protein fluorescent bright spots dropped to ∼50%, indicating that passive regulation based on membrane curvature may still exist in *P. aeruginosa*, albeit to a lesser degree.

Is the remarkable consistency in the spatial distribution of these two independent structural units an unintentional outcome or a hidden mystery? Given that Kulasekara et al. previously proposed that CheA and its phosphorylation state can cause c-di-GMP heterogeneity among individual *P. aeruginosa* cells through a specific PDE^51,52^, we sought to verify whether the phenotypic differences induced by CheY here are closely related to changes in the overall population level of c-di-GMP. We introduced a c-di-GMP monitoring system at the single-cell level and discovered that overexpression CheY substantially increases the intracellular c-di-GMP concentration. Previous studies generally believed that CheY-P, as a chemotactic regulatory protein, only influences flagellar motor switching. Here, we found that CheY-P, as a signaling molecule, can accomplish threshold-limited cross-pathway regulation. At low CheY-P concentrations, it is conservatively involved in motor switching regulation, as confirmed by previous studies. At high CheY-P concentrations, cell motility is suppressed, and c-di-GMP is triggered to express, though its specific mechanism needs further exploration. Unlike CheA, a structural protein of the chemotaxis complex, CheY is a protein that can shuttle in the cytoplasm and can also regulate the intracellular c-di-GMP level. This finding expands our understanding of the connection between the internal chemotaxis network and the c-di-GMP pathway. The intracellular environment is highly crowded^53^, and the proximal growth pattern shown in Fig. 5B has multiple physiological significances. On one hand, chemotaxis-related protein can precisely regulate cell motility with minimal synthesis costs. On the other hand, the chemotactic network and the flagellar motor are spatially integrated within the cell, ensuring their robust existence within the cell body and preventing mutual interference with collateral pathways.

Many bacteria possess multiple signal transduction pathways. The orderly operation of each pathway within the micrometer-scale space presents a fascinating scientific problem. Here, we utilized the well-known chemotaxis network as a starting point to shed light on this problem, providing inspiration for future in-depth understanding of related issues.

## Materials and methods

### Strains and cell culture

The strains and plasmids used in this study are listed in Table 1. The *Escherichia coli* TOP10 strain was used for standard genetic manipulations. A single-colony isolate was grown in 3 ml of LB broth (1% Bacto tryptone, 0.5% yeast extract and 1% NaCl) overnight to saturation on a rotary shaker (250 rpm) at 37°C. An aliquot was diluted 1:100 into 10 ml of LB broth and grew to exponential phase. Appropriate antibiotics were added if necessary to prevent plasmid loss: for *E. coli*, 15 μg/ml Gentamicin and 25 μg/ml Tetracycline; for *P. aeruginosa*, 30 μg/ml Gentamicin and 50 μg/ml Tetracycline. To induce protein expression, Isopropyl β-D-1-thiogalactopyranoside (IPTG, 1.0 mM) was added to strains with pME6032-derivative vectors, and arabinose (0.005%, 0.01%, 0.05%, 0.1%) was added to strains with pJN105-derivative vectors. 2 ml of cells were harvested by centrifugation at 2000×*g* for 2 min, washed twice in an equal volume of motility buffer (MB) [50 mM potassium phosphate, 15 μM EDTA, 0.15 M NaCl, 5 mM Mg^2+^ and 10 mM lactic acid (pH 7.0)]^54^, and resuspended in 5 ml MB for subsequent fluorescence observation.

**Table 1.**
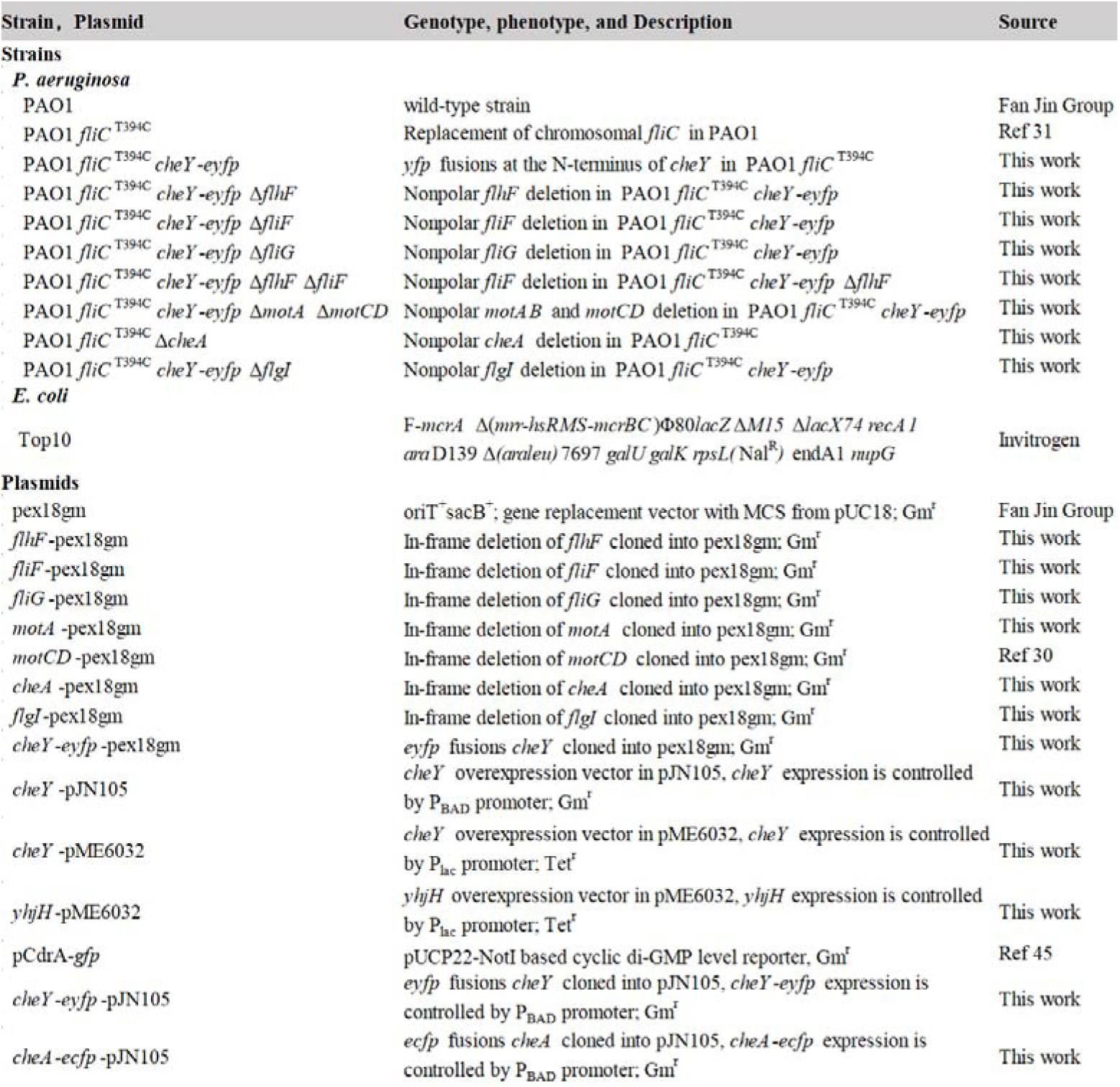
Strains and plasmids used in this study.

### Construction of in-frame deletion mutants

We used polymerase chain reaction (PCR) to generate ∼1000 bp DNA fragments with upstream (Up) and downstream (Dn) sequences flanking the target gene (including *flhF*, *fliF*, *motA* and *fliG*) to be deleted. The Up and Dn DNA sequences were fused with the linearized pex18gm vector using Gibson assembly^55^. The resulting vectors were transformed into *P. aeruginosa* by electroporation, and the desired knockout mutants were obtained by double selections on gentamicin plates (LB plates with 30 µg/ml gentamicin) and sucrose plates (NaCl-free LB plates with 15% sucrose) at 37°C^56^.

### Generation of chromosomal fusions of *yfp* to *P. aeruginosa* gene

To generate *eyfp* fusion at the N-terminus of *cheY*, we first used PCR to generate upstream and downstream DNA sequences flanking the *cheY* gene. A subsequent PCR reaction aimed to yield the *cheY-eyfp* fusion with a 3× glycine linker (GGCGGAGGA), with pVS88 serving as the source of the enhanced yellow fluorescent (*eyfp*). The three DNA sequences, along with the linearized vector, were fused to *cheY*-*eyfp*-pex18gm using Gibson assembly. Finally, gene replacement was used to transfer the fusion constructs into the *P. aeruginosa* chromosome by homologous recombination.

### Flagellar staining and fluorescence imaging

Flagellar filaments were labeled by following the protocol described previously^57^. Cells (1 ml of exponential-phase culture) were harvested by centrifugation at 2000×*g* for 10 min and washed twice in 1 ml MB. The final pellet was adjusted to a volume of ∼100 µl that concentrated the bacterial 10-fold. Alexa Fluor 568 maleimide (Invitrogen-Molecular Probes) was added to a final concentration of 20 μg/ml, and labeling was allowed to proceed for 30 min at room temperature with gyration at 80 rpm. Unused dye was then removed by washing the cells with MB three times, and the final pellet was resuspended in MB.

For fluorescence imaging, 50 μl of cells were added to the chamber (constructed with two double-sticky tapes, a glass slide, and a poly-L-Lysine coated coverslip), incubated for 3 min, and rinsed with 100 μl MB. The boundary of the chamber was then sealed with Apiezon vacuum grease. The chamber was placed on a Nikon Ti-E inverted fluorescence microscope with a 100× oil-immersion objective and a sCMOS camera (Primer95B, Photometrics). The flagellar filament and chemotaxis complex of cells were observed separately with corresponding filter-set, using a 200-ms exposure time. To quantify the fluorescence intensity of the receptor cluster, we took the intensity maximum point as the center and extract a 3×3 pixel matrix around it. The mean value of the elements in this matrix represents the final fluorescence intensity.

### Western Blotting

The total protein of *P. aeruginosa* strain cells in each treatment group was extracted and its concentration was measured with Bicinchoninic acid (BCA Protein Quantification Kit, Yeasen Biotechnology Co., Ltd) when the cells grew to exponential phase with same reduced turbidity (OD_600_ between 0.9 and 1.0). The exact amount of protein was subjected to SDS-PAGE electrophoresis, then transferred to a nitrocellulose membrane, which was blocked with 50 mg/mL skim milk in TBST buffer (20 mM Tris, 150 mM NaCl [pH7.4], 0.1%Tween 20) for 1h at room temperature. Since the N-terminal sequence of EYFP is almost identical to that of GFP, a rabbit anti-GFP antibody (1:2000, Yeasen Biotechnology Co., Ltd) and a Mouse anti-β-actin antibody (1:2000, Invitrogen) were added and incubated at 4°C overnight. The nitrocellulose membrane was gently washed with TBST for 20 min, three times. Subsequently, HRP labeled goat anti-rabbit IgG (1:4000, Abcam) and rabbit anti-mouse IgG (1:4000, Abcam) were added and incubated at room temperature for 1 h. The nitrocellulose membrane was again gently washed with TBST for 20 min, three times. Then, the membrane was developed using the ECL chemiluminescence detection system (Beyotime P0018FS) and visualized with the ChemiDoc XRS+ system (Bio-Rad). Three independent experiments were conducted for reproducibility.

### Monitoring c-di-GMP signal at the single-cell level

For measurements of c-di-GMP signaling, the validated reporter plasmid pCdrA-*gfp*, which produces green fluorescent protein (GFP) in response to an increase in c-di-GMP, was introduced into the experimental system. A 1000× diluted culture was inoculated into the apparatus described in the previous section and allowed to adhere to the surface for ∼10 min, with the slight difference that the coverslips were not coated with poly-L-Lysine to avoid interference from background fluorescence. Bacteria were imaged using a Nikon Ti-E inverted microscope, which was equipped with an EMCCD camera (DU897, iXon3, Andor Technology). We used a 100× oil-immersion objective. Bacteria were illuminated with a 488-nm laser (Sapphire 488−200 mW, Coherent), using a standard GFP filter sets in the fluorescence imaging system with an exposure time of 500 ms. To ensure the consistency of culture and observation conditions, the data of wild-type and *cheY*-pME6032 carried strains were collected at the same time. More than 200 independent individuals of each strain were used for data analysis.

Single-cell c-di-GMP concentration was characterized by calculating fluorescence intensity. The average fluorescence intensity of a single cell was calculated by dividing the total fluorescence intensity of a single cell by the cell volume. Bacterial were simplified to a hollow cylindrical trunk with hollow hemispherical caps at both ends. The cell volume can be calculated by taking the length of the short axis of the cell body as the diameter of the sphere and the length of the long axis as the sum of the height of the cylinder and the diameter of the sphere. The mean fluorescence intensity of the cell was obtained by subtracting the average background fluorescence intensity from the total fluorescence intensity of the cell, and then dividing by the cell volume.

## Author Contributions

J.Y. and R.Z. designed the work; Z.W., M.T. and S.F. performed the measurements with help from M.C.; Z.W and S.F analyzed the data; Z.W., J.Y. and R.Z. wrote the paper. Z.W., M.T. and S.F. contributed equally to this work.

## Conflicts of interest

The authors declare no competing interests.

## Acknowledgments

This work was supported by National Natural Science Foundation of China Grants (11925406, 12090053, 12304241, and 12304251), a Grant from the Ministry of Science and Technology of China (2019YFA0709303), Grants from the Natural Science Foundation of Shandong Province No. ZR2023QA111 and ZR2023QC168, a special Funding for High-level talents in the Medical and Health Industry in Jinan, a Grant from the Tai Shan Young Scholar Foundation of Shandong Province (tsqnz20231257) and the Science and Technology Development Program of Jinan Municipal Health Commission (2023-1-3).

---

**Fig. S1.**
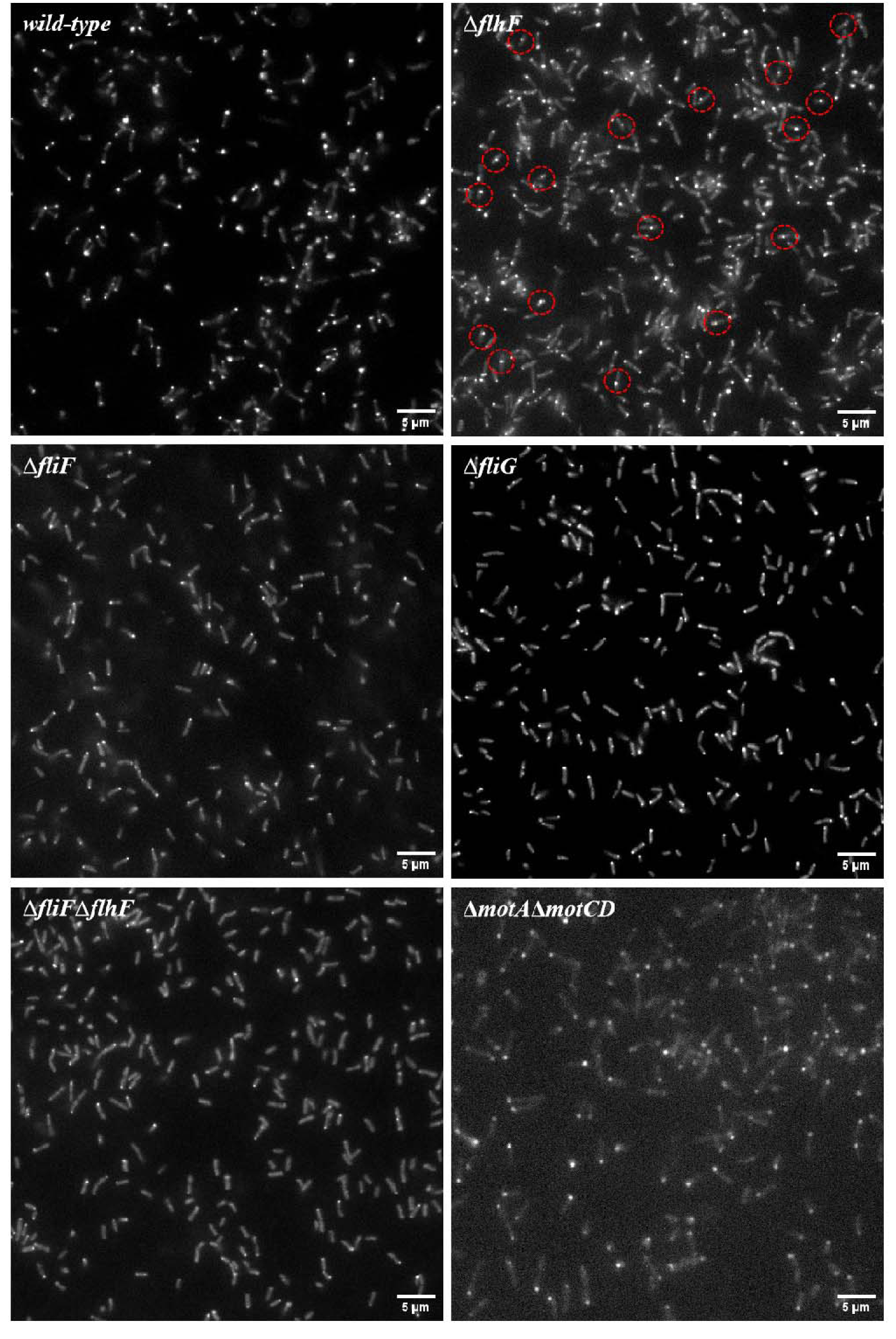
Distribution of chemotaxis complexes in multiple strains within representative large fields. Red dotted circles indicate cells where the chemotaxis complex is located at the mid-cell position.

**Fig. S2.**
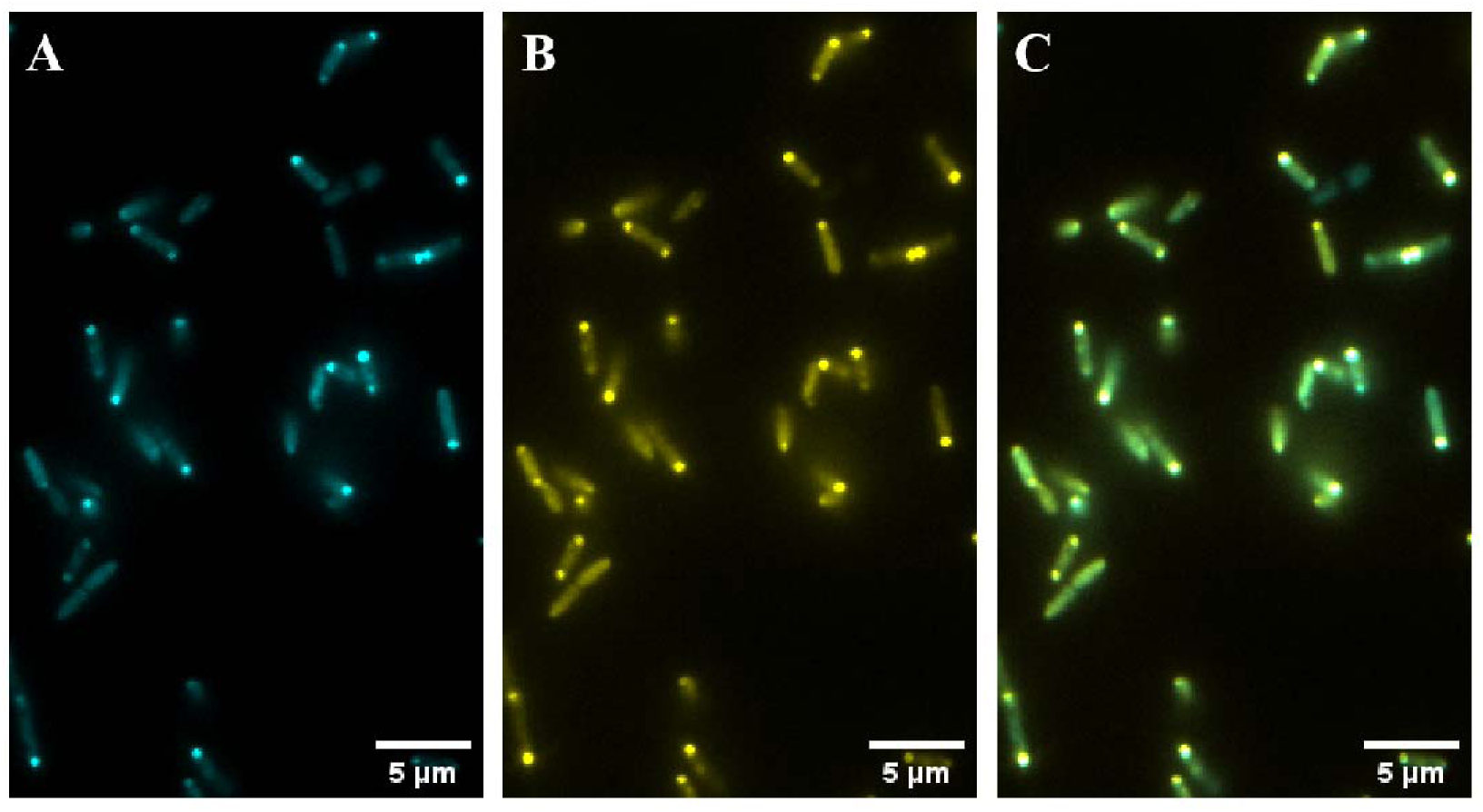
Co-localization of CheY-EYFP and CheA-ECFP in *P. aeruginosa*. CheA-ECFP is shown in blue (left), CheY-EYFP in yellow (center), and the merged image is shown in the right panel.

**Fig. S3.**
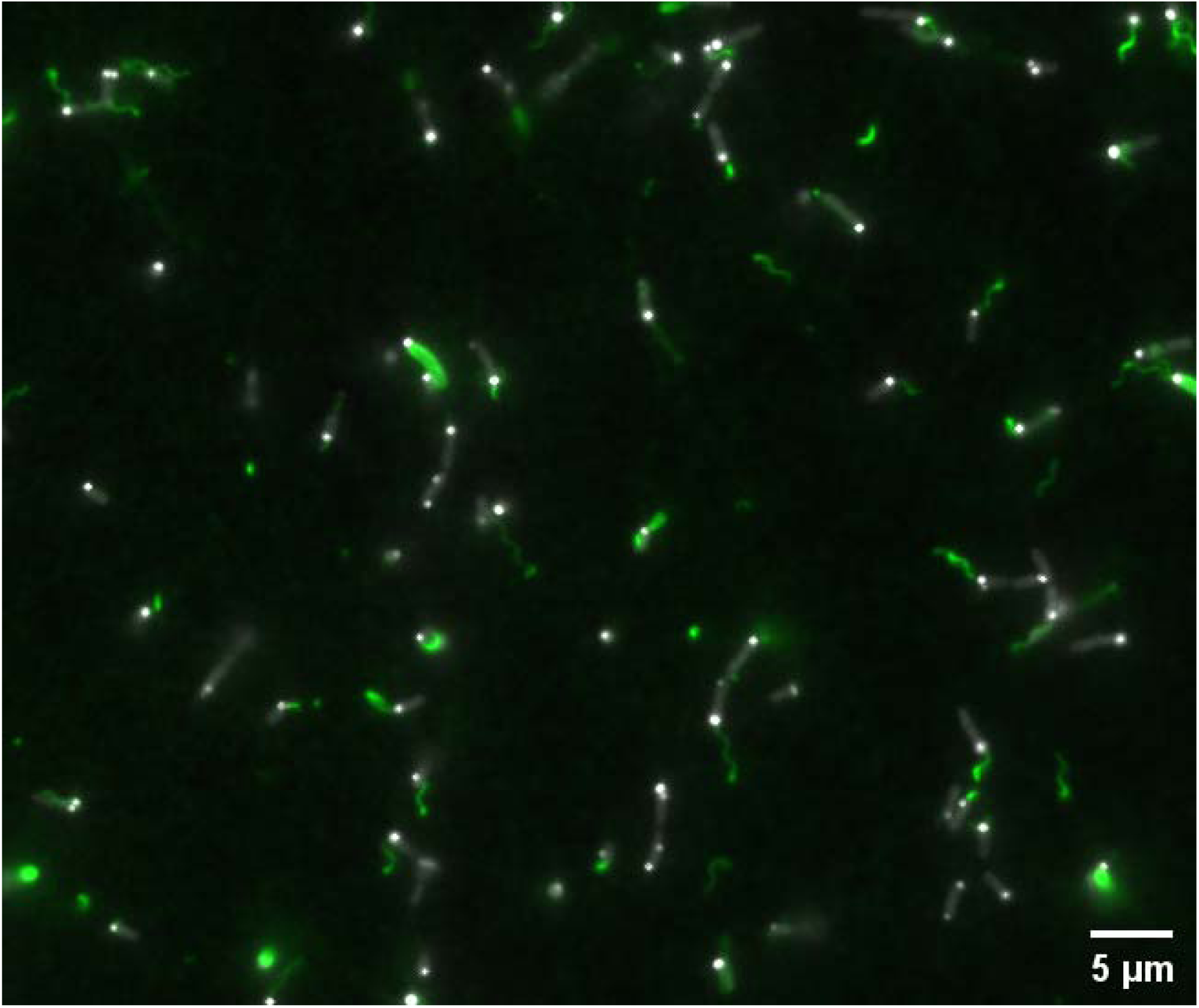
Simultaneous observation of the chemotaxis complex and flagellar filaments in the wild-type strain, shown in a representative large field.

**Fig. S4.**
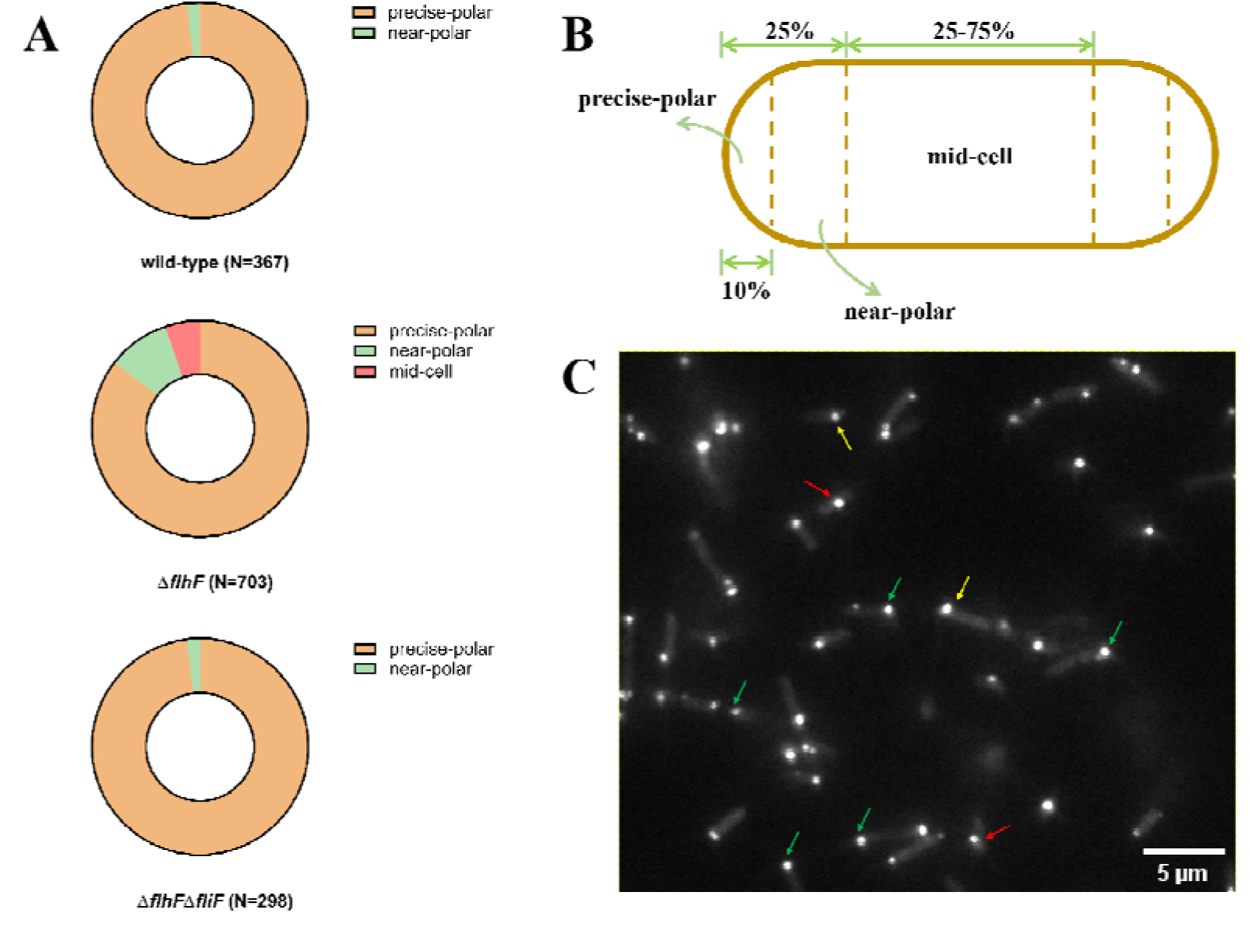
**A**. Distribution statistics of the chemotaxis complex in the wild-type strain and the *flhF* mutant. The distribution patterns are categorized into three types: precise-polar, near-polar, and mid-cell localization. In the wild-type strain, 98.1% of cells exhibit precise-polar localization of the chemotaxis complex, with the remainder showing near-polar distribution. In the *flhF* mutant, the proportion of cells with precise-polar localization decreases to 85.5%, and mid-cell localization appears in approximately 5% of cells. The *flhF-fliF* double mutant displays a distribution pattern similar to that of the wild-type strain. **B.** Schematic diagram of the classification of chemotactic complex locations. **C.** Cells with different chemotactic complex distribution patterns. Red arrows correspond to the mid-cell type, yellow arrows correspond to the near-polar type, and green arrows correspond to the precise-polar type.

**Fig. S5.**
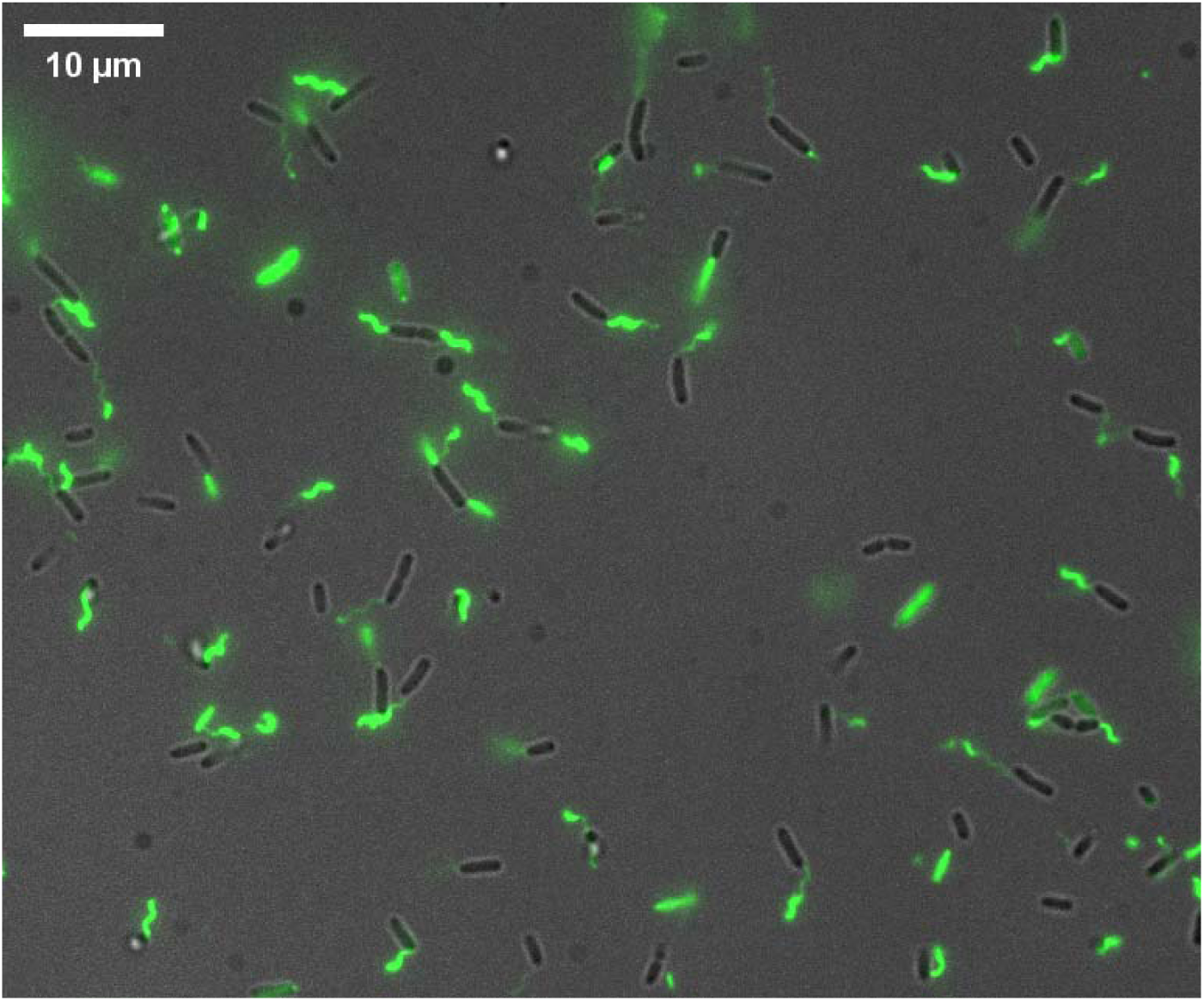
Knockout of *cheA* did not affect lagellar assembly efficiency of *P. aeruginosa*.

**Fig. S6.**
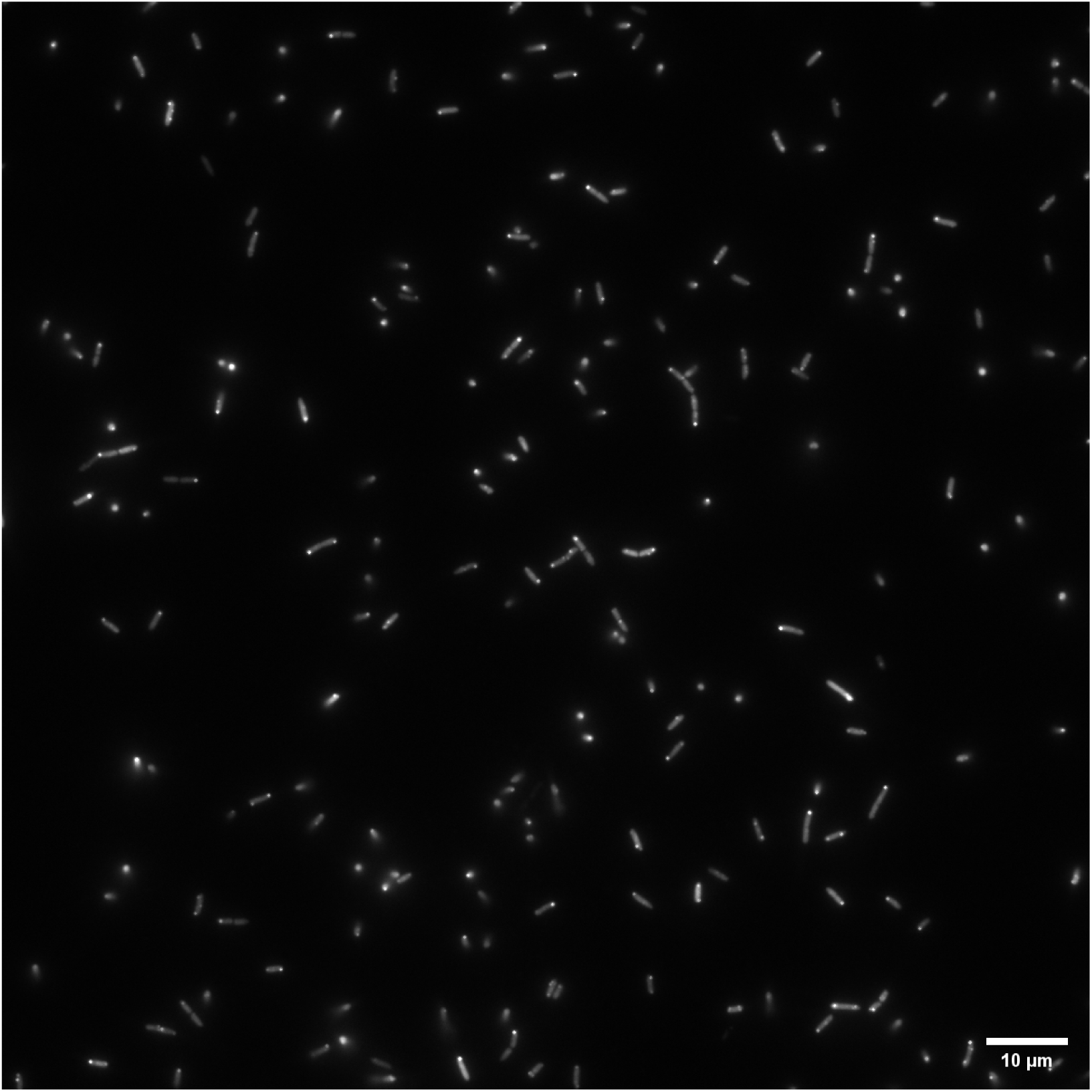
Localization of CheY-EYFP in the Δ*flgI* strain of *P. aeruginosa*. The phenotype is similar to that of the wild-type strain.

**Fig. S7.**
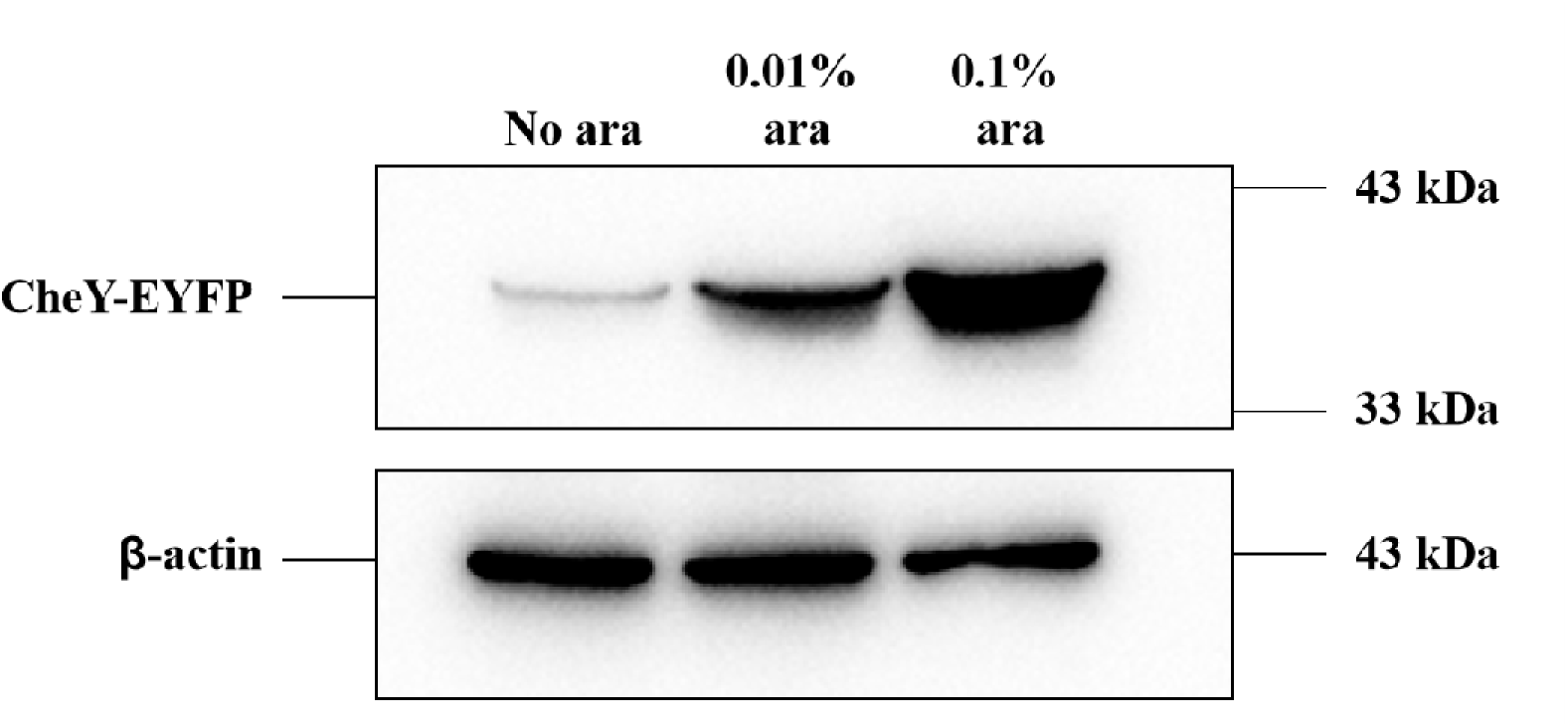
Intracellular CheY levels induced by different arabinose concentrations, measured by western blot analysis, showing clear differences across the concentration gradient. β-actin was used as the housekeeping protein.

**Fig. S8.**
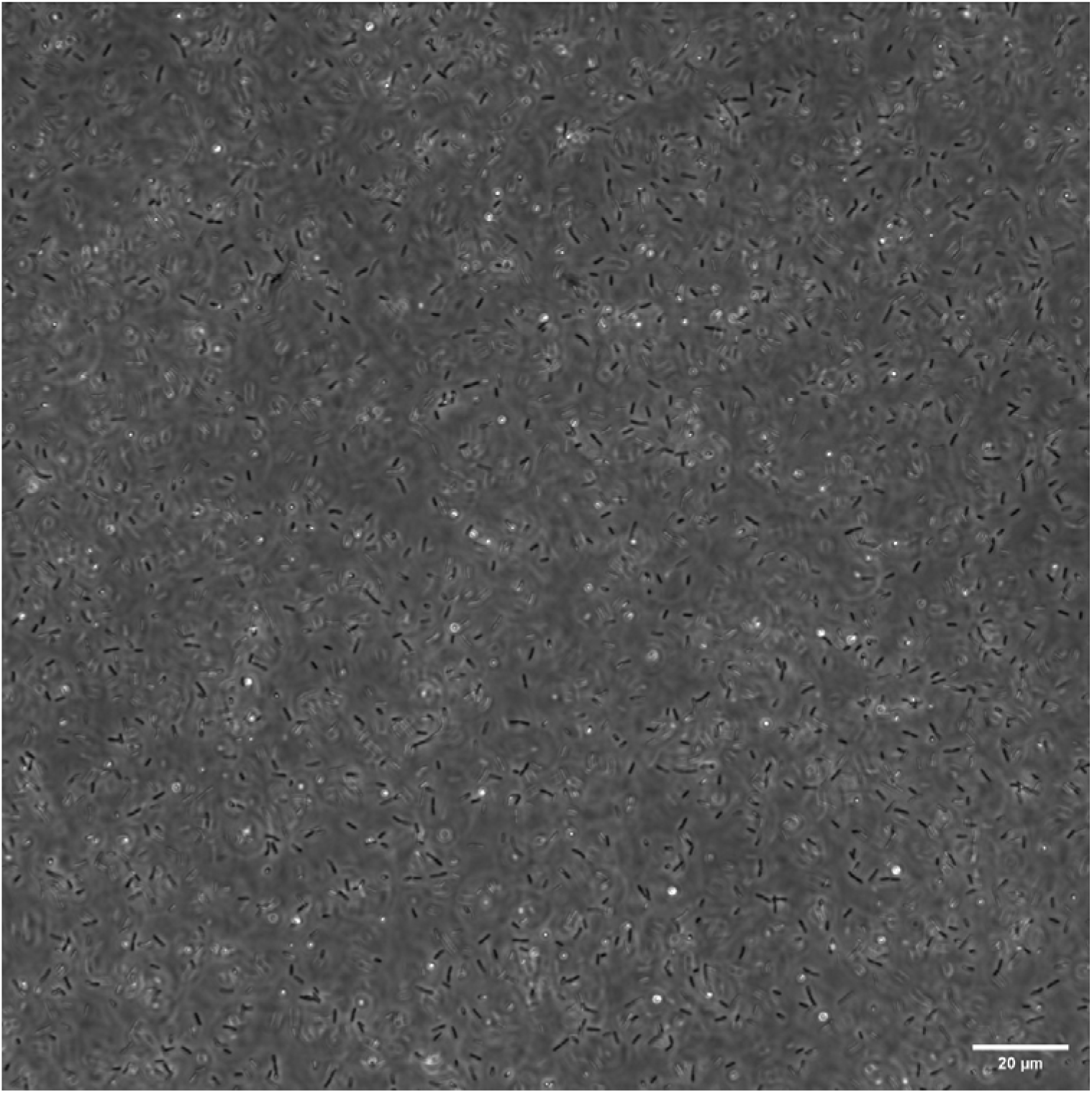
Co-overexpressing CheY and the phosphodiesterase (PDE) YhjH from *E. coli* mitigates cell aggregation caused by CheY overexpression in *P. aeruginosa*.

## Notes

### Competing Interest Statement

The authors have declared no competing interest.

### Summary of Updates

We clarified the categorization and spatial definitions of chemotaxis complex localization patterns, including the addition of a schematic diagram and representative examples (Fig. S4). We have expanded our discussion of the transcriptional regulatory differences between E. coli, Shewanella oneidensis, and P. aeruginosa. We now properly highlight that unlike E. coli, chemoreceptor expression in P. aeruginosa appears to be independent of flagellar assembly status.

## References

1 Halatek, J., Brauns, F. & Frey, E. Self-organization principles of intracellular pattern formation. Philos. Trans. R. Soc. Lond. B Biol. Sci. 373, 20170107 (2018).

2 Thanbichler, M. & Shapiro, L. Getting organized--how bacterial cells move proteins and DNA. Nat. Rev. Microbiol. 6, 28–40 (2008).

3 Loose, M., Kruse, K. & Schwille, P. Protein self-organization: lessons from the min system. Annu. Rev. Biophys. 40, 315–336 (2011).

4 Rothfield, L., Taghbalout, A. & Shih, Y. L. Spatial control of bacterial division-site placement. Nat. Rev. Microbiol. 3, 959–968 (2005).

5 Fay, A. & Glickman, M. S. An essential nonredundant role for mycobacterial DnaK in native protein folding. PLoS Genet. 10, e1004516 (2014).

6 Vaubourgeix, J. et al. Stressed mycobacteria use the chaperone ClpB to sequester irreversibly oxidized proteins asymmetrically within and between cells. Cell Host Microbe. 17, 178–190 (2015).

7 Wang, J., Brodmann, M. & Basler, M. Assembly and Subcellular Localization of Bacterial Type VI Secretion Systems. Annu. Rev. Microbiol. 73, 621–638 (2019).

8 Low, H. H. et al. Structure of a type IV secretion system. Nature 508, 550–553 (2014).

9 Briegel, A. et al. Bacterial chemoreceptor arrays are hexagonally packed trimers of receptor dimers networked by rings of kinase and coupling proteins. Proc. Natl. Acad. Sci. U. S. A. 109, 3766–3771 (2012).

10 Zhang, P., Khursigara, C. M., Hartnell, L. M. & Subramaniam, S. Direct visualization of Escherichia coli chemotaxis receptor arrays using cryo-electron microscopy. Proc. Natl. Acad. Sci. U. S. A. 104, 3777–3781 (2007).

11 Cluzel, P., Surette, M. & Leibler, S. An ultrasensitive bacterial motor revealed by monitoring signaling proteins in single cells. Science 287, 1652–1655 (2000).

12 Sourjik, V. & Berg, H. C. Binding of the Escherichia coli response regulator CheY to its target measured in vivo by fluorescence resonance energy transfer. Proc. Natl. Acad. Sci. U. S. A. 99, 12669–12674 (2002).

13 Welch, M., Oosawa, K., Aizawa, S. & Eisenbach, M. Phosphorylation-dependent binding of a signal molecule to the flagellar switch of bacteria. Proc. Natl. Acad. Sci. U. S. A. 90, 8787–8791 (1993).

14 Schuhmacher, J. S., Thormann, K. M. & Bange, G. How bacteria maintain location and number of flagella? FEMS Microbiol. Rev. 39, 812–822 (2015).

15 Jones, C. W. & Armitage, J. P. Positioning of bacterial chemoreceptors. Trends Microbiol. 23, 247–256 (2015).

16 Matilla, M. A., Martin-Mora, D., Gavira, J. A. & Krell, T. Pseudomonas aeruginosa as a Model To Study Chemosensory Pathway Signaling. Microbiol. Mol. Biol. Rev. 85, e00151–00120 (2021).

17 Guvener, Z. T., Tifrea, D. F. & Harwood, C. S. Two different Pseudomonas aeruginosa chemosensory signal transduction complexes localize to cell poles and form and remould in stationary phase. Mol. Microbiol. 61, 106–118 (2006).

18 Amako, K. & Umeda, A. Flagellation of Pseudomonas aeruginosa during the cell division cycle. Microbiol. Immunol. 26, 113–117 (1982).

19 Schniederberend, M., Abdurachim, K., Murray, T. S. & Kazmierczak, B. I. The GTPase activity of FlhF is dispensable for flagellar localization, but not motility, in Pseudomonas aeruginosa. J. Bacteriol. 195, 1051–1060 (2013).

20 Murray, T. S. & Kazmierczak, B. I. FlhF is required for swimming and swarming in Pseudomonas aeruginosa. J. Bacteriol. 188, 6995–7004 (2006).

21 Greenfield, D. et al. Self-organization of the Escherichia coli chemotaxis network imaged with super-resolution light microscopy. PLoS Biol. 7, e1000137 (2009).

22 Thiem, S. & Sourjik, V. Stochastic assembly of chemoreceptor clusters in Escherichia coli. Mol. Microbiol. 68, 1228–1236 (2008).

23 Strahl, H. et al. Transmembrane protein sorting driven by membrane curvature. Nat. Commun. 6, 8728 (2015).

24 Draper, W. & Liphardt, J. Origins of chemoreceptor curvature sorting in Escherichia coli. Nat. Commun. 8, 14838 (2017).

25 Koler, M., Peretz, E., Aditya, C., Shimizu, T. S. & Vaknin, A. Long-term positioning and polar preference of chemoreceptor clusters in E. coli. Nat. Commun. 9, 4444 (2018).

26 Huitema, E., Pritchard, S., Matteson, D., Radhakrishnan, S. K. & Viollier, P. H. Bacterial birth scar proteins mark future flagellum assembly site. Cell 124, 1025–1037 (2006).

27 Ringgaard, S., Schirner, K., Davis, B. M. & Waldor, M. K. A family of ParA-like ATPases promotes cell pole maturation by facilitating polar localization of chemotaxis proteins. Genes Dev. 25, 1544–1555 (2011).

28 Ringgaard, S. et al. ParP prevents dissociation of CheA from chemotactic signaling arrays and tethers them to a polar anchor. Proc. Natl. Acad. Sci. U. S. A. 111, E255–E264 (2014).

29 Sourjik, V. & Berg, H. C. Localization of components of the chemotaxis machinery of Escherichia coli using fluorescent protein fusions. Mol. Microbiol. 37, 740–751 (2000).

30 Wu, Z., Tian, M., Zhang, R. & Yuan, J. Dynamics of the Two Stator Systems in the Flagellar Motor of Pseudomonas aeruginosa Studied by a Bead Assay. Appl. Environ. Microbiol. 87, e0167421 (2021).

31 Tian, M., Wu, Z., Zhang, R. & Yuan, J. A new mode of swimming in singly flagellated Pseudomonas aeruginosa. Proc. Natl. Acad. Sci. U. S. A. 119, e2120508119 (2022).

32 Altegoer, F., Schuhmacher, J., Pausch, P. & Bange, G. From molecular evolution to biobricks and synthetic modules: a lesson by the bacterial flagellum. Biotechnol. Genet. Eng. Rev. 30, 49–64 (2014).

33 Kazmierczak, B. I. & Hendrixson, D. R. Spatial and numerical regulation of flagellar biosynthesis in polarly flagellated bacteria. Mol. Microbiol. 88, 655–663 (2013).

34 Minamino, T. & Imada, K. The bacterial flagellar motor and its structural diversity. Trends Microbiol. 23, 267–274 (2015).

35 Suzuki, H., Yonekura, K. & Namba, K. Structure of the rotor of the bacterial flagellar motor revealed by electron cryomicroscopy and single-particle image analysis. J. Mol. Biol. 337, 105–113 (2004).

36 Minamino, T., Imada, K. & Namba, K. Molecular motors of the bacterial flagella. Curr. Opin. Struct. Biol. 18, 693–701 (2008).

37 Levenson, R., Zhou, H. & Dahlquist, F. W. Structural insights into the interaction between the bacterial flagellar motor proteins FliF and FliG. Biochemistry 51, 5052–5060 (2012).

38 Hizukuri, Y., Yakushi, T., Kawagishi, I. & Homma, M. Role of the intramolecular disulfide bond in FlgI, the flagellar P-ring component of Escherichia coli. J. Bacteriol. 188, 4190–4197 (2006).

39 O’Connor, J. R., Kuwada, N. J., Huangyutitham, V., Wiggins, P. A. & Harwood, C. S. Surface sensing and lateral subcellular localization of WspA, the receptor in a chemosensory-like system leading to c-di-GMP production. Mol. Microbiol. 86, 720–729 (2012).

40 Terasawa, S. et al. Coordinated reversal of flagellar motors on a single Escherichia coli cell. Biophys. J. 100, 2193–2200 (2011).

41 Sagawa, T. et al. Single-cell E. coli response to an instantaneously applied chemotactic signal. Biophys. J. 107, 730–739 (2014).

42 Segall, J. E., Block, S. M. & Berg, H. C. Temporal comparisons in bacterial chemotaxis. Proc. Natl. Acad. Sci. U. S. A. 83, 8987–8991 (1986).

43 Qian, C., Wong, C. C., Swarup, S. & Chiam, K. H. Bacterial tethering analysis reveals a “run-reverse-turn” mechanism for Pseudomonas species motility. Appl. Environ. Microbiol. 79, 4734–4743 (2013).

44 Sourjik, V. Receptor clustering and signal processing in E. coli chemotaxis. Trends Microbiol. 12, 569–576 (2004).

45 Rybtke, M. T. et al. Fluorescence-based reporter for gauging cyclic di-GMP levels in Pseudomonas aeruginosa. Appl. Environ. Microbiol. 78, 5060–5069 (2012).

46 Vaknin, A. & Berg, H. C. Single-cell FRET imaging of phosphatase activity in the Escherichia coli chemotaxis system. Proc. Natl. Acad. Sci. U. S. A. 101, 17072–17077 (2004).

47 Dasgupta, N. et al. A four-tiered transcriptional regulatory circuit controls flagellar biogenesis in Pseudomonas aeruginosa. Mol. Microbiol. 50, 809–824 (2003).

48 Chilcott, G. S. & Hughes, K. T. Coupling of Flagellar Gene Expression to Flagellar Assembly in Salmonella enterica Serovar Typhimurium and Escherichia coli. Microbiol. Mol. Biol. Rev. 64, 694–708 (2000).

49 Beeby, M., Ferreira, J. L., Tripp, P., Albers, S.-V. & Mitchell, D. R. Propulsive nanomachines: the convergent evolution of archaella, flagella and cilia. FEMS Microbiol. Rev. 44, 253–304 (2020).

50 Kaplan, M. et al. The presence and absence of periplasmic rings in bacterial flagellar motors correlates with stator type. eLife 8, e43487 (2019).

51 Kulasekara, B. R. et al. c-di-GMP heterogeneity is generated by the chemotaxis machinery to regulate flagellar motility. eLife 2, e01402 (2013).

52. Kulasekara, B. R. Insight into a Mechanism Generating Cyclic di-GMP Heterogeneity in Pseudomonas aeruginosa PhD thesis, University of Washington, (2013).

53 Zhou, H., Rivas, G. & Minton, A. P. Macromolecular crowding and confinement: biochemical, biophysical, and potential physiological consequences. Annu. Rev. Biophys. 37, 375–397 (2008).

54 Cai, Q., Li, Z., Ouyang, Q., Luo, C. & Gordon, V. D. Singly Flagellated Pseudomonas aeruginosa Chemotaxes Efficiently by Unbiased Motor Regulation. mBio 7, e00013–00016 (2016).

55 Gibson, D. G. et al. Enzymatic assembly of DNA molecules up to several hundred kilobases. Nat. Methods 6, 343–345 (2009).

56 Hmelo, L. R. et al. Precision-engineering the Pseudomonas aeruginosa genome with two-step allelic exchange. Nat. Protoc. 10, 1820–1841 (2015).

57 Turner, L., Zhang, R., Darnton, N. C. & Berg, H. C. Visualization of Flagella during bacterial Swarming. J. Bacteriol. 192, 3259–3267 (2010).

